# Compromised chronic efficacy of a Glucokinase Activator AZD1656 in mouse models for common human *GCKR* variants

**DOI:** 10.1101/2024.07.24.604910

**Authors:** Brian E Ford, Shruti S Chachra, Ahmed Alshawi, Fiona Oakley, Rebecca J. Fairclough, David M Smith, Dina Tiniakos, Loranne Agius

## Abstract

Glucokinase activators (GKAs) have been developed as blood glucose lowering drugs for type 2 diabetes. Despite good short-term efficacy, several GKAs showed a decline in efficacy chronically during clinical trials. The underlying mechanisms remain incompletely understood. We tested the hypothesis that deficiency in the liver glucokinase regulatory protein (GKRP) as occurs with common human *GCKR* variants affects chronic GKA efficacy. We used a *Gckr*-P446L mouse model for the *GCKR* exonic rs1260326 (P446L) variant and the *Gckr-*del/wt mouse to model transcriptional deficiency to test for chronic efficacy of the GKA, AZD1656 in GKRP-deficient states. In the *Gckr*-P446L mouse, the blood glucose lowering efficacy of AZD1656 (3 mg/kg body wt) after 2 weeks was independent of genotype. However after 19 weeks, efficacy was maintained in wild-type but declined in the LL genotype, in conjunction with raised hepatic glucokinase activity and without raised liver lipids. Sustained blood glucose lowering efficacy in wild-type mice was associated with qualitatively similar but more modest changes in the liver transcriptome compared with the P446L genotype, consistent with GKA therapy representing a more modest glucokinase excess than the P446L genotype. Chronic treatment with AZD1656 in the *Gckr*-del/wt mouse was associated with raised liver triglyceride and hepatocyte microvesicular steatosis. The results show that in mouse models of liver GKRP deficiency in conjunction with functional liver glucokinase excess as occurs in association with common human *GCKR* variants, GKRP-deficiency predisposes to declining efficacy of the GKA in lowering blood glucose and to GKA induced elevation in liver lipids.

## 1. INTRODUCTION

With progression of type 2 diabetes (T2D) a combination of drugs targeting different mechanisms is used to compensate for declining pancreatic β-cell function or efficacy of individual therapies [1]. Glucokinase (GK), encoded by the *GCK* gene, was identified as a candidate target for T2D therapy because of its dual role in hepatic glucose metabolism and as glucose-sensor for insulin secretion and additionally because of evidence of compromised liver and islet GK in advanced T2D [2]. Several classes of GK activator drugs (GKAs) were developed with good acute efficacy in lowering blood glucose [3]. However, few have been approved for T2D therapy or are still in development [4-9]. Key challenges encountered were risks of hypoglycaemia by excessive insulin secretion and a chronic decline in glucose-lowering efficacy [10-12]. Proposed hypotheses for declining efficacy, include β-cell failure consequent to over-activation of GK in islet β-cells and hepatic steatosis by over-activation of GK in liver [13,14].

In preclinical studies sustained GKA efficacy in lowering blood glucose for up to 8 wk had been reported for various rodent models of insulin resistance [13,15-17] and for up to 40 wk in models of genetic *Gck* deficiency [18,19], which are characterized by stable hyperglycaemia similar to human *GCK* haplo-insufficiency (*GCK*-Maturity onset diabetes of the young) [20]. However, loss in GKA glycaemic efficacy was found in rodent models of spontaneous diabetes such as the Goto-Kakizaki rat [21] and *db/db* mouse [22-24], which are characterized by dysregulated insulin secretion and raised pancreatic islet GK activity [25,26]. In the *db/db* mouse, β-cell failure consequent to GK excess in islet β-cells was demonstrated by the protective role of *Gck* haplo-insufficiency on diabetes progression [27] and by the restoration of glucose sensing in *db/db* pancreatic islets by inhibiting glucose phosphorylation with mannoheptulose [28]. Whether raised liver GK activity contributes to declining GKA efficacy on blood glucose has not been reported.

Liver GK which accounts for ∼99% of total GK [2] is regulated at the transcriptional level by insulin and glucagon and on a rapid scale in response to changes in blood glucose or fructose by adaptive binding to the glucokinase regulatory protein GKRP, encoded by the *GCKR* gene [29,30]. GKRP is located in the hepatocyte nucleus and sequesters GK in the nucleus in an inactive state at basal blood glucose enabling its rapid release and translocation to the cytoplasm in response to elevated glucose or fructose in the absorptive state [29]. Common human *GCKR* gene variants comprising a missense variant (rs1260326, P446L) with compromised binding to GK [31,32] and intronic variants with frequencies ranging from 0.15 to 0.5 in different populations associate with a wide range of metabolic traits [33,34]. A mouse model for the missense variant (*Gckr*:P446L) replicates several of the metabolic traits associated with the *GCKR* variants in human populations [35]. In this study we used the *Gckr*:P446L mouse and a model of transcriptional *Gckr* deficiency (*Gckr*^del/wt^) to test the hypothesis that compromised GKRP function or expression affects the response to chronic GKA treatment. Using AZD1656, a GKA with an established safety record [36,37] that has been extensively trialled in man [6,11,12], we show that compromised GKRP function (*Gckr*:P446L) or expression (*Gckr*^del/wt^) affect chronic blood glucose lowering efficacy and lipid accumulation in liver.

## 2. MATERIALS AND METHODS

### 2.1. Animals and ethics statement

Mice were bred and maintained in compliance with the Animals Scientific Procedures Act, 1986. All procedures and animal studies were approved by Newcastle University Animal Welfare Ethics Review Board in compliance with UK Home Office Regulations under license PC1B783F4, and in line with the ARRIVE guidelines. Two mouse lines on the C57BL/6N background were used: *Gckr*^P446L^ (P446L; GCKR-P446L-EM1-B6N) [35] and the *Gckr-DEL* line (GCKR-DEL1262-EMI-B6N: C57BL/6N-Gckr<em1(IMPC)H>/H), through the International Mouse Phenotyping Consortium (IMPC:www.mousephenotype.org/data/genes/MGI:1096345). Mice were bred as heterozygous mating pairs. For GCKR-P446L-EM1-B6N the genotyping was via a custom Taqman SNP genotyping assay (#4332075, Assay No. ANU66PU) from ThermoFisher (Paisley, UK). For GCKR-DEL1262-EM1-B6N, genotyping, PCR primers (For: TGTCCACTCTCTACAGCGAC; Rev: TAAGGGTGCAAACTCAGCCG) were used in a three-step protocol (1 cycle of 98C for 5 min; 40 cycles of (98C denaturation (5s), 61.2C annealing (5s), 72C extension (20s)), 1 cycle of (72C for 1 min, 4C hold)). Products were run on a 2% agarose gel (100V, 30 min) and visualized with ethidium bromide. Expected product sizes are ∼1880 bp for wild-type, 618 bp for homozygous delete (del/del) and both products for heterozygous (del/wt) mice.

### 2.2. AZD1656 and animal diet

AZD1656 [16] and was provided by AstraZeneca (Cambridge, UK). The high-fat diet (HFD; #824018) from Special Diet Services (Whitham, Essex, UK) contained 45% fat (lard); 20% protein 35% carbohydrate (rice starch, sucrose, cellulose) by energy. The high-fat high-sugar diet (HFHSD) comprised the same HFD but with 10% glucose and 5% fructose in the drinking water.

### 2.3. Chronic mouse studies

#### 2.3.1. *Gckr:*P446L study with AZD1656 for 20 wk on HFHSD

Six groups of male mice of PP, PL or LL genotypes (aged 8 wk) received either HFD pellets (#824018) without AZD1656 (-) or HFD pellets with AZD1656 (+ 3 mg / kg body wt) for 20 wk (PP-, PP+, PL-, PL+, LL-, LL+ n=14,16,11,12,16,15) [16,19]. The drinking water contained 10% glucose and 5% fructose. Blood was sampled at the start of the study and after 2 wk and 19 wk in the *ad libitum* fed state for analysis of glucose and plasma insulin. Mice were caged by mixed genotype. An insulin tolerance test (ITT) was performed after 12 wk and glucose tolerance test after 18 wk. At the end of the study a terminal blood sample was taken for clinical chemistry analysis and tissues were harvested after terminal anaesthesia.

#### 2.3.2. *Gckr*^del/wt^ study with AZD1656 for 16 wk on HFD

Four groups of male mice of *Gckr*^wt/wt^ or *Gckr*^del/wt^ genotypes (aged 18wk) received either HFD (#824018) without AZD1656 (-) or HFD with AZD1656 (+ 3 mg / kg body wt) for 16 wk (wt/wt-, wt/wt+, del/wt-, del/wt+; n= 12,12,9,12) [16,19]. Blood was sampled at the start of the study and after 4-wk and 14 wk in the *ad libitum* fed state for analysis of glucose and plasma insulin. A glucose tolerance test was performed after 8 wk and insulin tolerance test after 12 wk. At the end of the study a terminal blood sample was taken for clinical chemistry analysis and tissues were harvested after terminal anaesthesia.

### 2.4. Blood and tissue analysis

Blood glucose was determined by tail vein sampling using an AccuChek glucose meter from Roche (West Sussex, UK) and plasma insulin by Mercodia (Uppsala, Sweden) ELISA (# 10-1247-01). For insulin tolerance tests, food was withdrawn 5h before blood sampling followed by intraperitoneal insulin (Henry Schein Animal Health, Dumfries, UK) injection (Actrapid, 0.75 Units/ kg body wt) and tail vein blood sampling for glucose measurement at the times indicated. For oral glucose tolerance tests food was withdrawn for 2h before gavage (2g glucose / kg body wt) [16,19] with tail vein blood sampling for glucose measurement at the times indicated. Plasma lipids, enzyme activities and inorganic phosphate analysis was performed at Mary Lyon Centre Pathology (AU680 analyzer). Liver lipids were extracted with chloroform / methanol and the lipid extract in the chloroform phase was dried, reconstituted in isopropanol and assayed for triglyceride (WAKO triglyceride kit) and total cholesterol (CAY10007640) using kits from Alphalaboratories (Hampshire, UK) and Cambridge Biosciences (Cambridge, UK) respectively.

### 2.5. Histopathology

Liver and pancreas tissue blocks were fixed in 4% formalin / phosphate-buffered saline and sectioned (at 4 μm thickness). Liver tissue sections were stained with haematoxylin and eosin (H&E) and Sirius Red Fast Green (SFRG) for histological scoring by a liver pathologist (DT) blinded to genotype or treatment, for steatosis, microsteatosis, hepatocyte ballooning and fibrosis according to the NASH Clinical Research Network scoring system [38]. Pancreas sections were immunostained for insulin and glucagon by the Royal Victoria Infirmary Pathology service. For assessment of islet area, the fractional insulin versus haematoxylin staining was determined by Aperio software and is expressed as % insulin / haematoxylin staining.

### 2.6. Immunohistochemical staining for GK and GKRP

Liver and pancreas tissue sections were stained for GK (SC-7908, 1:100 (Santa Cruz Biotechnology, Heidleberg, Germany) or PT-159629, 1:800 (ProteinTech, Manchester, UK)) and liver sections for GKRP (SC-6340 at 1:400 or 1:800 dilution) using the Roche Diagnostics (West Sussex, UK) Ventana automated system (BenchMark XT) with anti-rabbit (GK) or anti-goat (GKRP) horse-radish peroxidase conjugated anti-IgG. Slides were scanned (x20) with Aperio Image Software (Leica Biosystems). Scans were annotated and quantitative image analysis of the annotated regions was performed using Aperio Brightfield Image Analysis Software (Leica Biosystems) for nuclear H-scores (N-H) and cytoplasmic H-scores (C-H) for liver and C-H scores for pancreas. Data exported comprised the total number of nuclei and surrounding cytoplasmic regions within 3 intensity thresholds. N-H and C-H scores correspond to the equation ([% <u>w</u>eak] + [2 x % <u>m</u>oderate] + [3 x % <u>h</u>igh] within the range of 0 to 300. N-H thresholds (200,160,120 for w,m,h) are of higher intensity than C-H thresholds (210,180,150 for w,m,h) to take into account the higher nuclear intensity staining for GKRP and GK compared with cytoplasmic GK staining and accordingly N-H/C-H ratios are an underestimate of relative nuclear localization. Pancreatic islet mass was determined from the sum of % weak + % medium + % strong. Relative GK staining was determined from the H-score of [2 x % moderate] + [3 x % high] with thresholds of 175 and 100 as % weak was at background.

### 2.7. Immunoblotting for GK and GKRP

Liver tissue was homogenized in buffer containing 100mM Tris-HCl, pH 7.4, 100mM NaCl, 25mM NaF, 2mM EDTA, 0.1% Triton-X100, 1 mM benzamidine and protease inhibitors (P8340 from Sigma (Poole, UK)). Protein was assayed on the 14,000 x *g* supernatant and samples (15-45 μg protein) were resolved by SDS-PAGE on 10% SDS or 4-12% SDS gels, electrotransferred to PVDF membrane and blotted for GKRP (rabbit AZ680 from AstraZeneca (Cambridge, UK)) or GK (Proteintech PT15629) with horse radish peroxidase conjugated anti-rabbit IgG and developed by Enhanced Chemiluminescence. Densitometry was quantified by ImageJ [35].

### 2.8. Liver total glucokinase activity

Liver tissue was homogenized in 15 volumes of buffer (50mM HEPES, 100mM KCl, 1mM EDTA, 2.5 mM DTT, pH 7.5) and centrifuged at 100,000 x *g* (45 min). The lipid infranatant and pellet were discarded and hexokinase kinetic assays were performed at 100mM and 0.5mM glucose in medium containing 50mM HEPES, 100 mM KCl, 2mM MgCl_2_, 5mM ATP-Mg^2+^, 0.5mM NAD, 2.5mM DTT an 8 U/ml glucose 6-phosphate dehydrogenase (*Leuconostoc mesenteroides*) (G8404, Merck (Gillingham, UK)). GK activity was determined from the difference in hexokinase activity at high and low glucose [35].

### 2.9. RNA analysis by RT-qPCR and RNA Sequencing

Liver RNA was extracted in Trizol (ThermoFisher, Paisley, UK) and cDNA was synthesized using MMLV (M1705) and amplified with GoTaq qPCR Master Mix (A6002) from Promega (Hampshire, UK) with the primers listed previously [35]. Relative gene expression was determined using the delta-delta Cycle threshold and expressed as a ratio to a house keeping gene (18S or *RplpO*). For RNA-sequencing liver RNA was extracted and purified using Trizol followed by Qiagen (Manchester, UK) RNeasy (#74104) with on-column DNase digestion (#79254). RNA-sequencing was at the Newcastle University Genomics Core after quality assessment (RIN > 6). All samples had quality scores > 35, no adapter contamination and did not require trimming. Transcript counts were against reference sequences (M26, GRCm39) from Gencode (gencodegenes.org/mouse/). Differential gene expression was determined with DSeq2 in R via RStudio. Differentially expressed genes (DEGs) were defined as a log2-fold change > 30% and an adjusted P-value (FDR, false discovery rate) < 0.05. Pairwise comparison by genotype or by AZD1656 treatment was performed to determine DEGs.

### 2.10. Statistical analysis

Data is presented as mean ± SEM. Statistical analysis was by Prism 9.5 Software (Graphpad Inc) or SPSS (v27). Data was tested for outliers and normality and comparisons between groups of continuous variables was assessed by the two-tail t-test for parametric data and or by Mann Whitney for non-parametric, unless otherwise stated. For histopathology categorical scores data was analysed by Chi-square test. Statistical significance is set at P < 0.05 unless otherwise indicated.

## 3. RESULTS

### 3.1. AZD1656 enhanced liver GK activity in the *Gckr*-P446L mouse on HFHSD

*Gckr*-P446L knock-in mice are outwardly indistinguishable from wild-type with no difference in body weight (Figure 1A) on regular diet (RD) or after 20 wk on high-fat high-sugar diet (HFHSD). They showed no difference in liver *Gckr* mRNA levels by genotype on either diet (Figure 1B). Liver *Gck* mRNA was not different by genotype on RD but was lower in LL mice on HFHSD (Figure 1B). At the protein level, GKRP was below detection limits in LL mice on RD by immunostaining (Figure 1C) and immunoblotting (Figure 1D,E). GK immunoactivity was 30% lower in LL mice (Figure 1F) and the GK-to-GKRP protein ratio was higher on RD and HFHSD (Figure 1G). By immunostaining GKRP was predominantly nuclear in wild-type and was faintly detectable in LL mice on HFHSD (Figure 1H). There was no significant effect of AZD1656 treatment on *Gckr* and *Gck* mRNA or immunoactive protein (Figure 1E,F).

**Figure 1:**
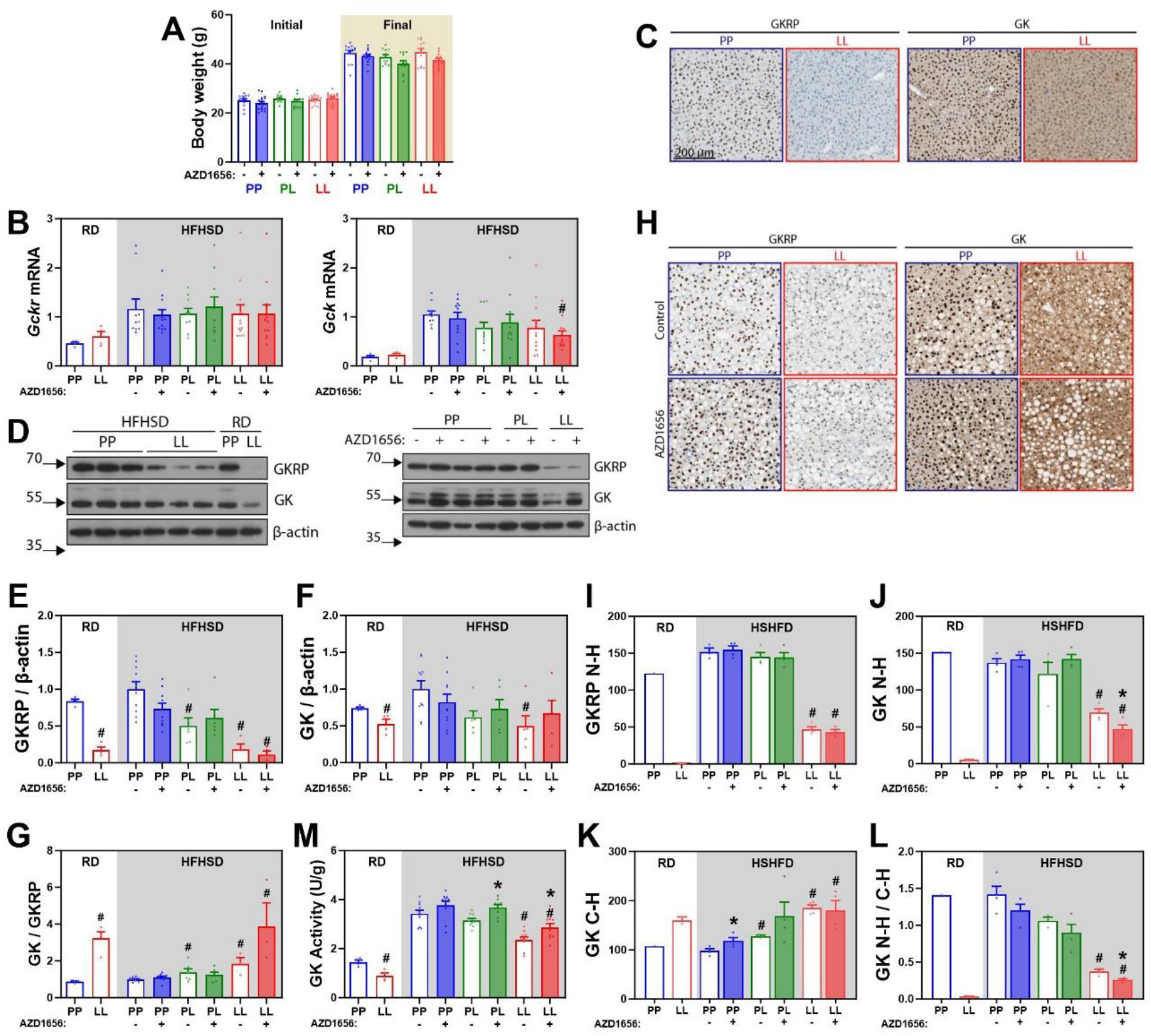
AZD1656 treatment increased liver GK activity in the P446L mouse. *Gckr*-P446L mice of PP, PL and LL genotypes were fed on a high-fat high-sugar diet (HFHSD) without (-, open bar) or with (+, filled bar) 3 mg/kg AZD1656 for 20 wk. A) Body weight at the start and end of the 20 wk study. B) Lack of effect of AZD1656 on *Gckr* mRNA and *Gck* mRNA (ratio to 18S). C) Liver GKRP and GK Immunostaining in 8 wk old P446L mice on regular diet (RD): showing no nuclear GKRP staining and cytoplasmic GK distribution in LL mice. D) Representative immunoblots for GKRP and GK in P446L mice on either RD or after 20 wk on HFHSD. E) GKRP immunoactivity on RD or after 20 wk HFHSD study showing lower protein in LL mice. F) GK immunoactivity on RD or after 20 wk HFHSD study showing lower protein in LL mice. G) GK/GKRP immunoactivity on RD or after 20 wk HFHSD showing higher GK/GKRP in LL mice. H) Representative GKRP and GK immunostaining after 20 wk HFHSD. I-L) Nuclear (N-H) and cytoplasmic (C-H) H-scores for GKRP and GK on RD and at the end of the 20 wk study (n=4) I) Lower GKRP N-H scores in LL mice. J) AZD1656 decreased GK N-H scores in LL mice. K) AZD1656 increased GK C-H scores in PP (wild-type) mice. L) AZD1656 decreased GK nuclear distribution (N-H/C-H) in LL mice. M) AZD1656 increased liver GK activity in PL and LL mice. \**P < 0*.*05* effect of AZD1656; #*P < 0*.*05* effect of genotype.

To test for differences in GK nuclear-cytoplasmic distribution we determined nuclear (N-H) and cytoplasmic (C-H) H-scores, a semi-quantitative estimate of staining intensity (Figure 1I-L). For GKRP, N-H scores were lower by LL genotype with no effect of AZD1656 treatment (Figure 1I). For GK, AZD1656 increased cytoplasmic GK distribution in PP and LL genotypes as shown by the higher C-H scores in PP mice and lower N-H scores and N-H/C-H ratios in LL mice (Figure 1J-L). Total GK enzyme activity was 40% lower in LL mice on RD (Figure 1M). GK activity increased > 2-fold after 20 wk on HFHSD and was 30% lower in LL mice (Figure 1M). AZD1656 treatment significantly increased total GK activity by 21% and 17% in LL and PL genotypes with a smaller (9%) trend in wild-type.

To test for a direct effect of the HFD on liver GK activity, mice of PP and PL genotypes were fed on RD or HFD for 2 wk. The mice on the HFD had higher fat pad weight and higher blood triglyceride and cholesterol (Figure 2). Liver GK activity was lower in PL mice than in wild-type mice (by 21%) but it was increased in PL mice by 24% (P<0.04) by the HFD. Cumulatively, AZD1656 increased total GK activity (∼ 20%) in PL and LL genotypes and it increased the cytoplasmic-to-nuclear GK distribution.

**Figure 2:**
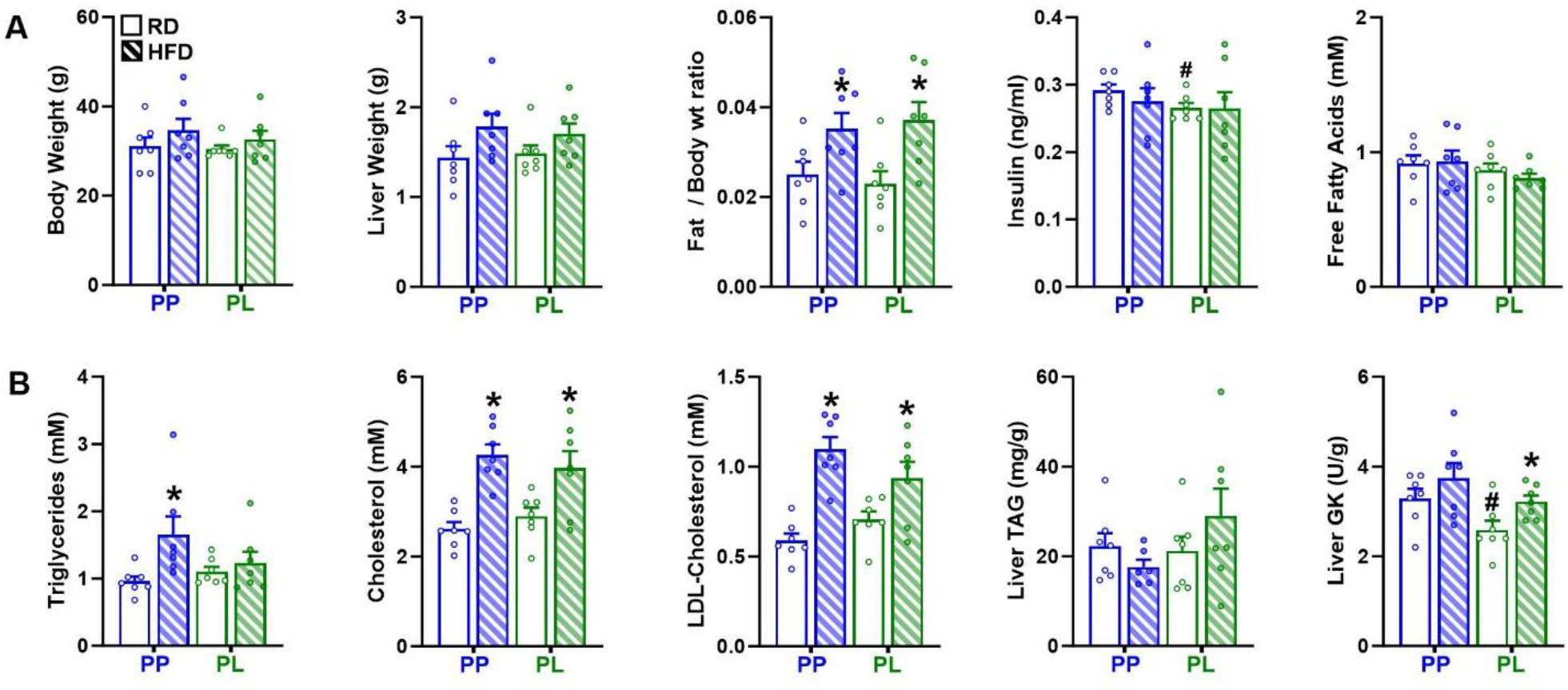
Effects of a high-fat diet on blood lipids and liver GK activity. P446L mice of the PP or PL genotypes (n=7,7) were fed on either regular diet (RD, open bar) or the high-fat diet (HFD, hashed) for 2 wk. A) Increase in epididymal fat pad mass on the HFD. B) Increase in blood triglyceride and cholesterol and increase in liver GK activity in PL mice on the HFD. \**P < 0*.*05* effect of diet; #*P < 0*.*05* effect of genotype.

### 3.2. Sustained blood glucose lowering by AZD1656 in wild-type but not in LL mice

There was no difference in blood glucose by P446L genotype on RD (Figure 3A, 0 wk). Treatment with AZD1656 for 2 wk lowered blood glucose by ∼3 mM (Figure 3B) in all genotypes (PP 3.1 ± 0.32; PL 3.1 ± 0.49; LL 3.5 ± 0.58 mM), and after 19 wk the glucose lowering was sustained in wild-type (3.2 ± 0.28 mM) but was attenuated in the LL genotype (1.7 ± 0.66 mM, P< 0.002) and blood glucose was significantly higher after 19 wk compared to 2 wk (6.5 ± 0.43 vs 4.8 ± 0.28 mM) in AZD1656-treated LL-mice (Figure 3A). Plasma insulin was not different by genotype or AZD1656 treatment after 2 wk but was lower by PL and LL genotypes after 19 wk (Figure 3C). Glucose tolerance determined after 18 wk was not different by genotype in either untreated or AZD1656 treated groups (Figure 3D). There was no difference by genotype in pancreatic islet area estimated from positive insulin staining (Figure 3E,F). Pancreatic islet GK immunostaining estimated from a modified C-H score showed no difference by genotype but higher staining by AZD1656 treatment in wild-type (Figure 3G). Cumulatively, this shows a decline in blood glucose lowering at 19 wk in LL mice but not in wild-type mice in the absence of a decline in islet GK staining.

**Figure 3:**
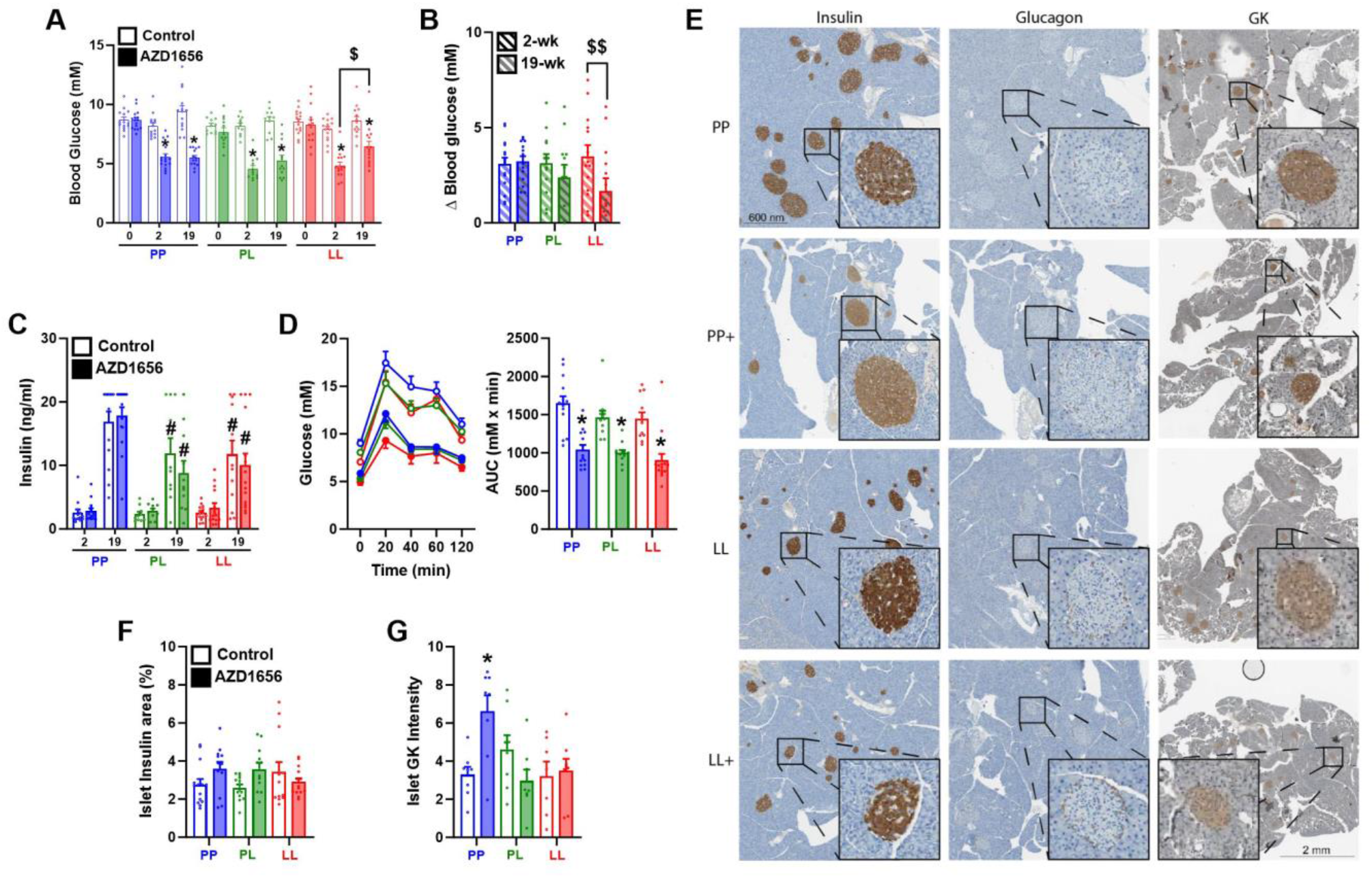
Chronic attenuation in blood glucose lowering by AZD1656 in the P446L mouse despite sustained islet GK staining. Experimental details were as in Figure 1. A) Similar blood glucose by genotype at the start (0 wk) of the 20 wk study and sustained blood glucose lowering by AZD1656 (3mg/kg) after 19 wk compared to 2 wk in wild-type (PP) but not in LL mice. B) Difference (delta Δ) in blood glucose by AZD1656 treatment after 2 wk (white stripe) and 19 wk (grey stripe) relative to start of study (0 wk) showing lower efficacy at 19 wk in LL mice. C) Lack of effect of AZD1656 on plasma insulin. D) Glucose tolerance test after 2h food withdrawal determined at 18wk: glucose excursion and area under the curve (AUC). E) Representative immunostaining of pancreas for insulin, glucagon and GK after 20 wk HFHSD -/+ AZD1656 in PP and LL mice. F) Pancreatic islet % area determined from insulin staining. G) Islet GK-staining intensity showing higher intensity by AZD1656 treatment in wild-type, PP mice. \**P < 0*.*005* effect of AZD1656; #*P < 0*.*05* effect of genotype; $*P < 0*.*01; $$ P< 0*.*005* 19 wk *versus* 2 wk.

### 3.3. No effect of chronic AZD1656 treatment on blood or liver lipids in LL mice

There was no effect of AZD1656 treatment on plasma triglycerides or free fatty acids but total cholesterol and LDL-cholesterol were higher by AZD1656 treatment in wild-type (11% and 18%, respectively) and they were higher by LL genotype (21% and 32%, respectively) but not further increased by AZD1656 (Figure 4A). There was no effect of AZD1656 on liver lipids or liver enzymes in plasma but inorganic phosphate was lower by LL genotype but unchanged by AZD1656 treatment (Figure 4B). Liver histopathology scores showed that there was no difference by genotype or AZD1656 treatment for hepatocyte steatosis, microvesicular steatosis, inflammation or fibrosis but lipogranuloma scores were higher in wild-type by AZD1656 treatment (Figure 4C).

**Figure 4:**
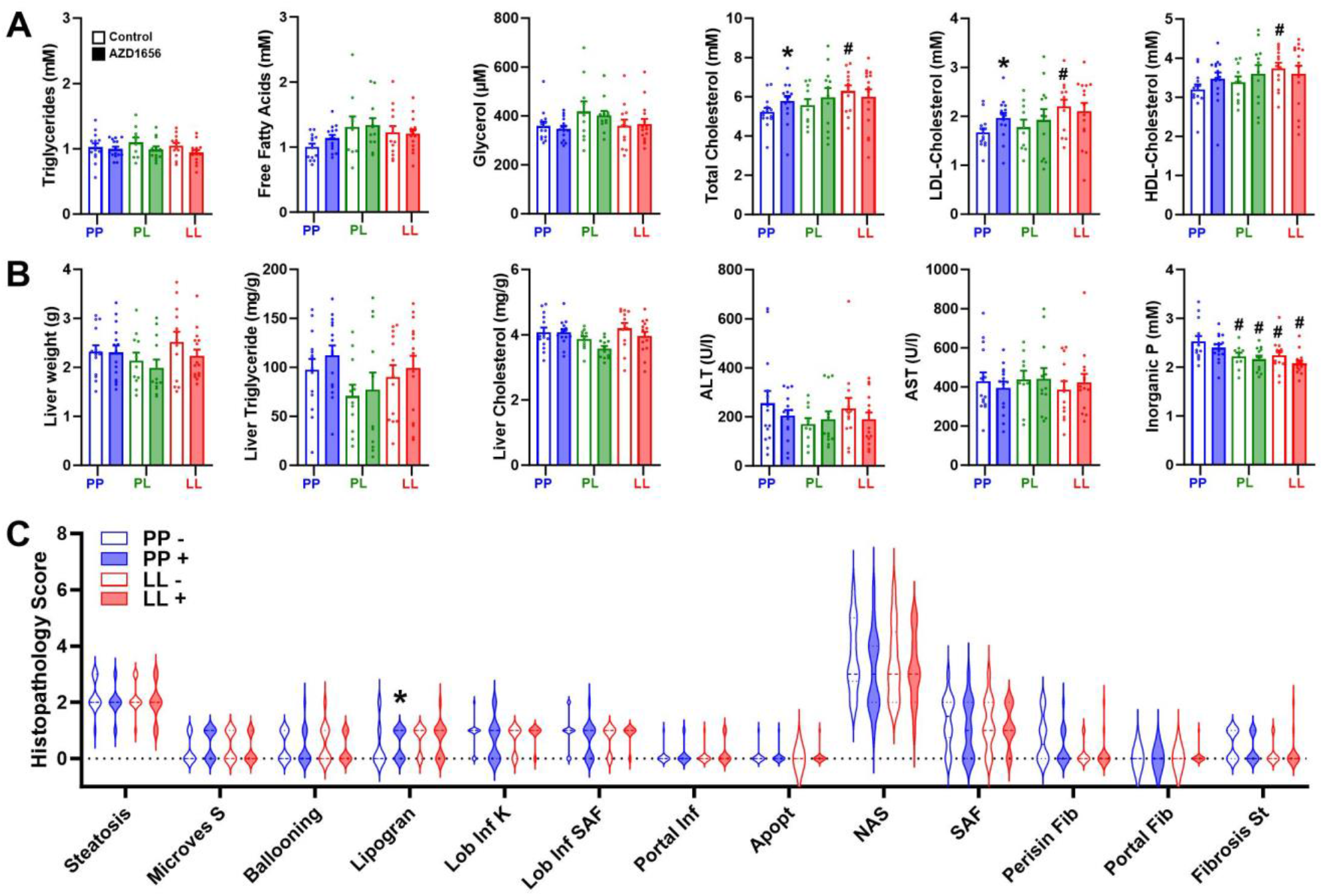
Lack of effect of AZD1656 treatment on blood or liver triglyceride and cholesterol in P446L mice after 20 wk. Experimental details were as in Figure:1. A) Blood triglycerides and cholesterol after 20 wk on HFHSD showing increased cholesterol (total and LDL) by LL genotype and by AZD1656 treatment in PP but not LL mice. B) Lack of effect of AZD1656 on liver triglyceride and cholesterol and liver enzymes (ALT, alanine aminotransferase; AST, aspartate aminotransferase) in plasma. Decrease in plasma inorganic phosphate by genotype but not by AZD1656. C) Lack of effect of AZD1656 on liver steatosis and fibrosis scores in PP and LL mice. \**P < 0*.*05* effect of AZD1656; #*P < 0*.*05* effect of genotype.

### 3.4. Modest effects of AZD1656 on the liver transcriptome compared with the P446L genotype

To assess the hepatic adaptations to chronic AZD1656 treatment we used RNA-sequencing and compared the liver transcriptome differences by P446L genotype (LL versus PP) with the AZD1656 treatment in PP or LL mice after 20 wk. There were 150 differentially expressed genes (DEGs, *adjusted-P*<0.05) by genotype in untreated mice (PP^-^/LL^-^) but far less by AZD1656 treatment in PP mice (26 DEGs; PP^-/+^) or LL mice (13 DEGs; LL^-/+^, Figure 5A). For the DEGs by genotype (PP^-^/LL^-^) Enrichment Gene Ontology analysis (EnrichGO) identified response to unfolded protein, organic acid transmembrane transport and the cholesterol biosynthesis pathway amongst top enriched pathways (not shown) and Ingenuity Pathway Analysis (IPA) identified protein ubiquitination, oxidative stress response, unfolded protein response, cholesterol biosynthesis, mitochondrial dysfunction and autophagy amongst top enriched pathways (Figure 5B). Comparison of DEGs by GKA treatment in PP^-/+^ or LL^-/+^ mice, with the DEGs by genotype (PP^-^/LL^-^) showed that for wild-type more than half (14 of 26 DEGs) were in common (and directionality) with the DEGs by genotype (PP^-^/LL^-^).

**Figure 5:**
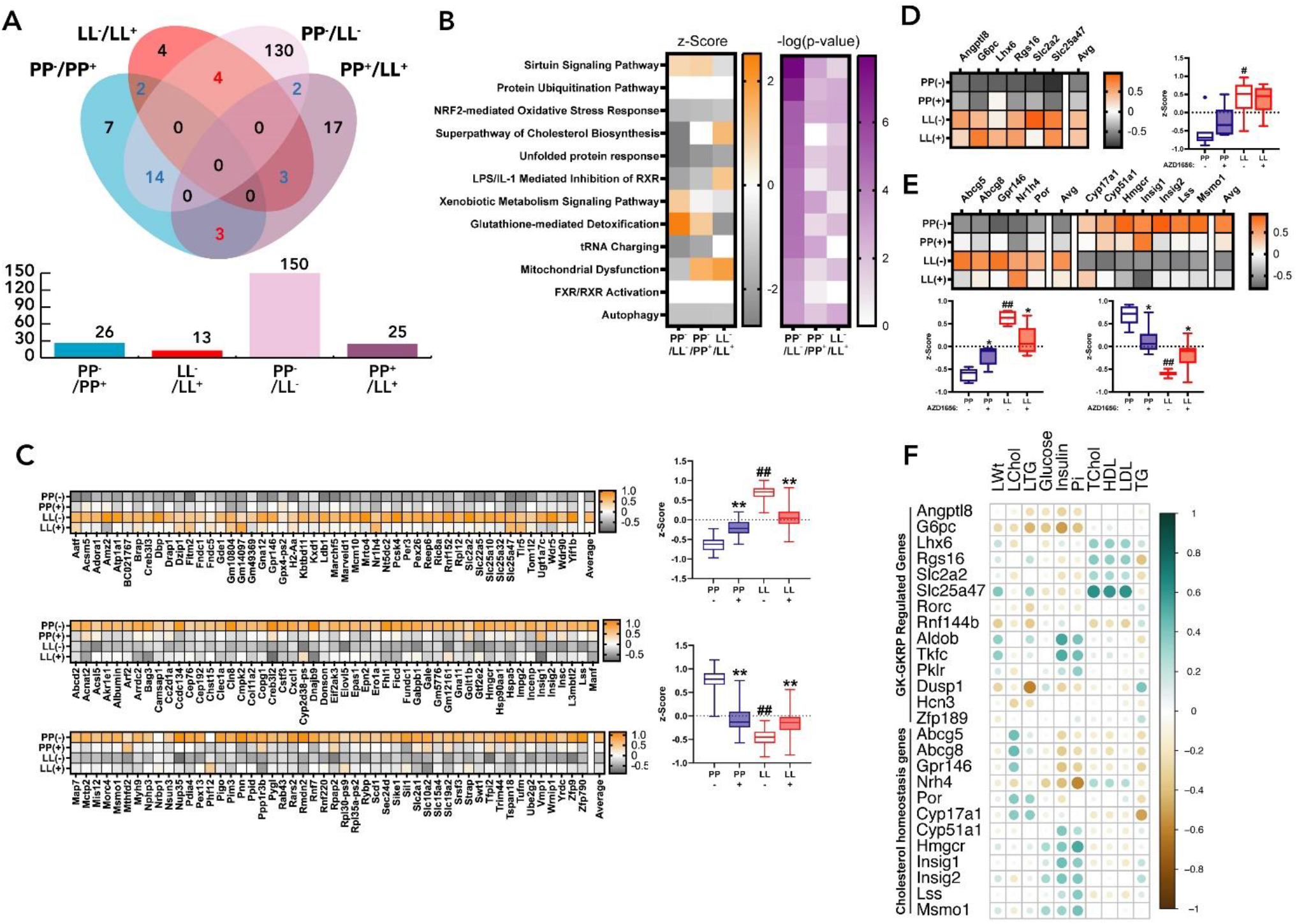
Liver transcriptome analysis in P446L mice after 20 wk on HFHSD -/+ AZD1656: lesser impact of AZD1656 treatment compared with P446L genotype. A) Venn diagram for differentially expressed genes (DEGs adjusted P < 0.05) by AZD1656 treatment: PP-vs PP+ (26 DEGs) and LL-vs LL+ (13 DEGs); or by genotype PP-vs LL-(150 DEGs), PP+ vs LL+ (25 DEGs). Blue, common genes similar direction; red, common genes opposite direction. B) Ingenuity Pathway Analysis: showing top enriched pathways by genotype (PP-vs LL-) or by AZD1656 treatment (PP-/PP+ or LL-/LL+) showing Z-scores for pathway flux and –log (P-value). C-E) Genes that are differentially expressed by PP-vs LL-genotype show smaller but directionally similar changes by AZD1656 treatment in PP mice. Z-scores for gene counts and median and range of genes shown. \*\**P<0*.*001*, \**P<0*.*02* effect of AZD1656; ##*P<0*.*0001* genotype effect, one-way ANOVA. C) Z-scores for DEGs by PP>LL genotype (150 DEGs adjusted P < 0.05) showing relative gene counts for the 4 groups of mice (PP-, PP+, LL-, LL+) of the upregulated DEGs (n=49, upper panel) and downregulated DEGs (n=101). D) Z-scores of gene counts of GK and GKRP-responsive genes that are significantly elevated by LL genotype. E) Z-scores of gene counts of cholesterol homeostasis linked genes that are significantly up-regulated or down-regulated by LL genotype. F) Correlation of phenotypic traits (liver weight, liver cholesterol, liver triglyceride, blood glucose and insulin and blood triglycerides and blood cholesterol Total, LDL< HDL) with corresponding gene counts of GK-GKRP responsive genes identified in [35] and cholesterol homeostasis linked genes.

To further compare the adaptations to chronic AZD1656 with those by P446L genotype the Z-scores of the gene counts were analysed for the 150 DEGs by genotype (PP^-^/LL^-^) for the four groups of mice (PP and LL untreated or AZD1656-treated). This analysis showed that many of the DEGs that were up-regulated or down-regulated by LL genotype (PP^-^/LL^-^) showed similar trends by AZD1656 treatment in the PP mice (PP^-^ >PP^+^), but to a lesser extent than by PP>LL genotype (Figure 5C). Similar trends were identified for genes linked to functional GK excess (Figure 5D) or cholesterol homeostasis (Figure 5E). Previous work identified 14 genes that are induced by short-term GK overexpression and counter-regulated by human and mouse GKRP (446P) in hepatocytes [35]. Of these, 6 genes were up-regulated by LL genotype (Figure 5D) and included *Slc25a47, Rgs16* and *Lhx6* which correlated positively with blood cholesterol as well as *G6pc* which correlated negatively with liver triglyceride, blood glucose, insulin and inorganic phosphate (Figure 5F). The other GK-GKRP responsive genes [35] which were not differentially expressed by LL genotype included *Aldob* and *Tkfc* which correlated positively with inorganic phosphate and insulin (Figure 5F). Of the cholesterol-linked genes (Figure 5E), the upregulated genes by LL genotype correlated positively with liver cholesterol and negatively with insulin whereas genes downregulated by LL genotype correlated positively with insulin and inorganic phosphate (Figure 5F). Cumulatively this shows directionally similar changes in a range of transcripts for AZD1656 treatment of wild-type but not LL mice as for the DEGs by P446L genotype.

### 3.5. AZD1656 treatment enhanced hepatic steatosis in *Gckr*^del/wt^ mice on HFD

As a second model of GKRP deficiency we used the *Gckr*^del/wt^ mouse on HFD. On RD in the *Gckr*^del/del^ genotype, there was no detectable nuclear GKRP immunostaining whereas the *Gckr*^del/wt^ had modest GKRP deficiency (Figure 6A). Treatment with AZD1656 for 16 wk on HFD had no effect on body weight (Figure 6B) or on GKRP and GK N-H-scores (immunostaining intensity) or GK N-H/C-H scores which were ∼30% lower than in *Gckr*^wt/wt^ (Figure 6C,D). By immunoblotting, *Gckr*^del/wt^ mice had more modest GKRP deficiency than *Gckr*-P446L mice (30% vs 80% lower in LL genotype, Figure 6E). In wild-type mice, AZD1656 had no effect on GKRP immunoactivity but increased the GK-to-GKRP protein ratio (Figure 6F). In *Gckr*^del/wt^ mice, AZD1656 likewise did not affect GKRP protein levels but increased GK and GK/ GKRP immunoreactivity (Figure 6G). Liver GK enzyme activity was lower (30%) in *Gckr*^del/wt^ than in wild-type and was increased (19%, P < 0.05) by AZD1656 in *Gckr*^del/wt^ with a smaller trend in wild-type (Figure 6H). Cumulatively, this model of modest GKRP and GK deficiency (30%), shows an increase in liver total GK enzyme activity and in GK-to-GKRP protein ratio by AZD1656 treatment.

**Figure 6:**
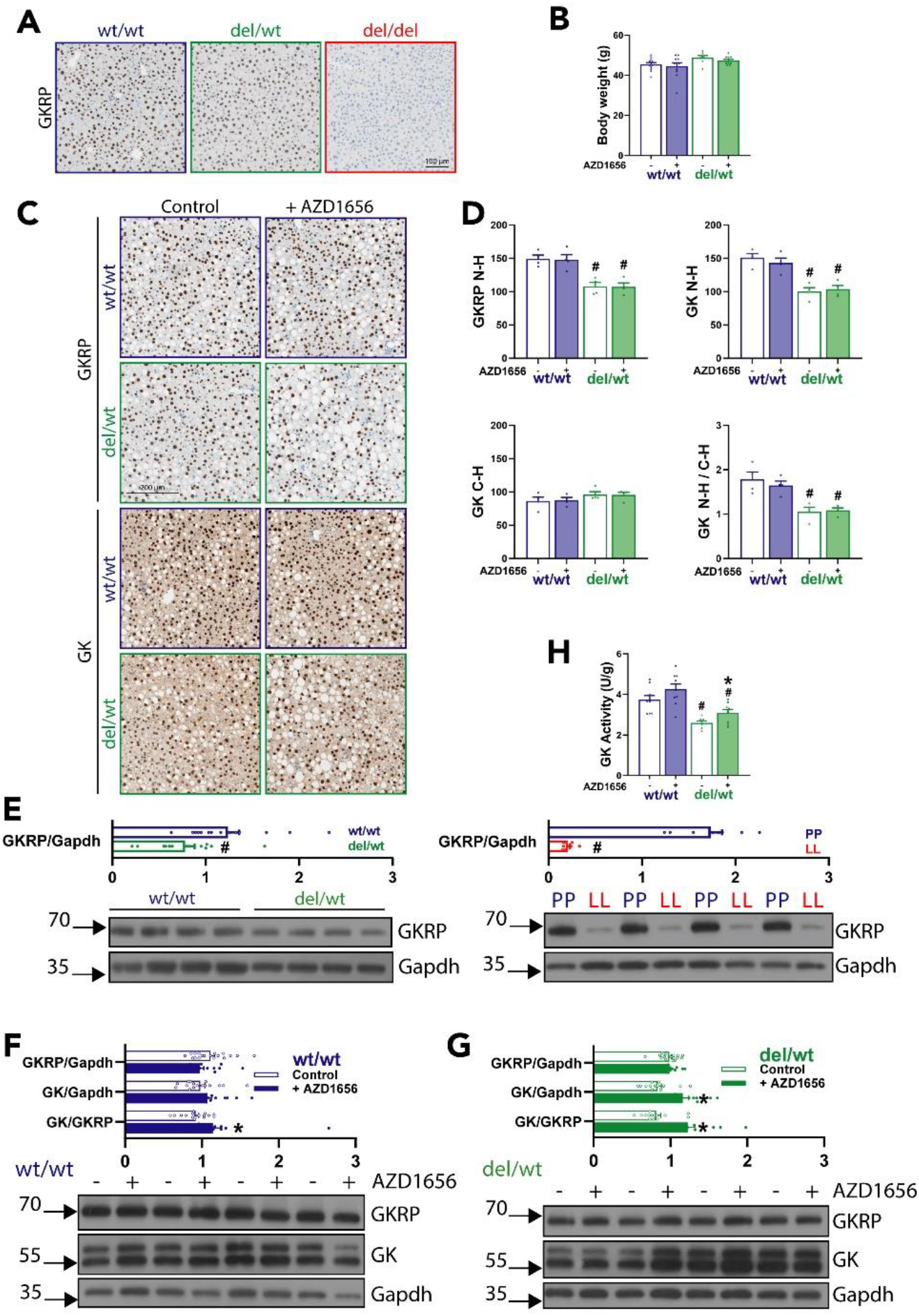
AZD1656 increased liver GK activity in the *Gckr*^*del/wt*^ mouse. *Gckr*^wt/wt^ and *Gckr*^del/wt^ were fed on a HFD without (open bar) or with (shaded bar) 3 mg/kg AZD1656 for 16 wk. A) Representative liver GKRP immunostaining of *Gckr*^wt/wt^, *Gckr*^del/wt^ and *Gckr*^del/del^ genotypes on regular diet. B) Body weight after 16 wk on HFD -/+ AZD1656. C) Representative liver immunostaining for GKRP and GK at the end of the study on HFD -/+ 3 AZD1656. D) GKRP N-H scores; GK N-H scores, GK C-H scores and GK N-H/C-H ratio showing lower GKRP and GK nuclear staining and greater cytoplasmic GK distribution by del/wt genotype. E) GKRP-immunoactivity by Western blot showing more modest GKRP deficiency in del/wt genotype (30%) than in LL (80%) genotype. F) GKRP and GK immunoactivity after 16 wk study in *Gckr*^wt/wt^ showing an increase in GK / GKRP immunoactivity ratio by AZD1656 treatment. G) GKRP and GK immunoactivity after 16 wk study in *Gckr*^del/wt^ mice showing an increase in GK and GK / GKRP immunoactivity ratio by AZD1656 treatment. H) Lower liver GK activity in the *Gckr*^del/wt^ mice and increase by AZD1656 treatment. \**P < 0*.*05* effect of AZD1656; #*P < 0*.*05* effect of genotype.

Blood glucose and insulin in the *Gckr*^del/wt^ mouse were similar to wild-type at the start of the study but plasma insulin was higher in *Gckr*^del/wt^ after 4 wk and 14 wk (Figure 7A,B). AZD1656 treatment lowered blood glucose after 4 wk and 14 wk by 32% and 27%, in wild-type and by 16% and 18%, in *Gckr*^del/wt^ mice without significant effect on plasma insulin (Figure 7A,B). There was no significant difference in glucose tolerance by AZD165 treatment but area under the curve was higher by *Gckr*^del/wt^ genotype in AZD1656-treated mice (Figure 7C).

**Figure 7:**
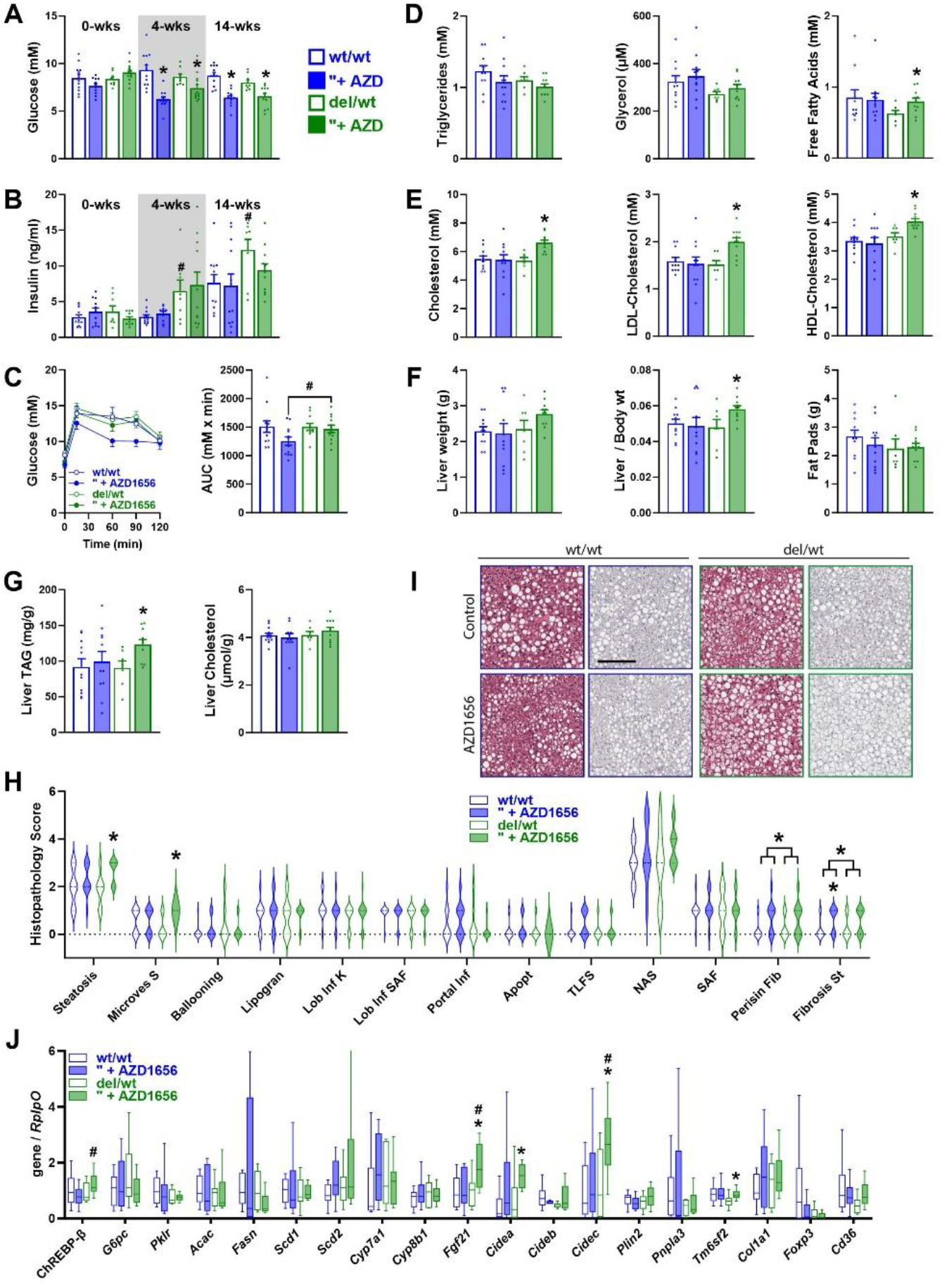
AZD1656 increases liver steatosis in the Gckr^del/wt^ mouse. *Gckr*^wt/wt^ and *Gckr*^del/wt^ were fed on a HFD without (open bar) or with (shaded) 3 mg/kg AZD1656 for 16 wk. A) Lowering of blood glucose by AZD1656 after 4 wk and 14 wk in *Gckr*^wt/wt^ and *Gckr*^del/wt^ mice. B) Higher plasma insulin by *Gckr*^del/wt^ genotype at 4 wk and 14 wk. C) Impaired glucose tolerance in AZD1656-treated *Gckr*^del/wt^ mice. D) Increased plasma FFA by AZD1656 treatment in *Gckr*^del/wt^ *mice*. E) Increased plasma cholesterol by AZD1656 treatment in *Gckr*^del/wt^ mice. F) Increased liver / body weight ratio by AZD1656 treatment in *Gckr*^del/wt^ mice. G) Increased liver triglyceride by AZD1656 treatment in *Gckr*^del/wt^ *mice*. H) Higher hepatocyte steatosis, microvesicular steatosis and fibrosis scores by AZD1656 treatment in *Gckr*^del/wt^ mice. I) Representative haematoxylin and eosin (H&E) and Sirius red fast green (SRFG) images for histopathology scores, scale bar 200μm. J) Liver mRNA levels (RT-qPCR) for the genes indicated expressed as a ratio to *RplpO* showing increased expression of *Fgf21, Cidea* and *Cidec* by AZD1656 treatment in the *Gckr*^del/wt^ mice: n= 10 and 11 for *Gckr*^wt/wt^ (-/+) and n= 6 and 10 for *Gckr*^del/wt^ (-/+) \**P < 0*.*05* effect of AZD1656; #*P < 0*.*05* effect of genotype.

AZD1656 treatment did not affect plasma triglyceride but raised free fatty acids (27%), total cholesterol, LDL and HDL cholesterol (24%, 31%, 16%, respectively) in *Gckr*^del/wt^ (Figure 7D,E). AZD1656 increased the liver / body wt ratio and liver triglyceride but not cholesterol (Figure 7F,G). Histopathological scoring for steatosis and fibrosis showed that hepatocyte steatosis was evident in most mice but AZD1656 treated *Gckr*^del/wt^ mice had higher scores for hepatocyte steatosis (P < 0.052) and microvesicular steatosis (P< 0.002). Scores for perisinusoidal fibrosis were modestly increased by AZD1656 in the combined genotypes (P < 0.05) and fibrosis sub-stage scores were higher by AZD1656 treatment in wild-type (P< 0.03) and in the combined genotypes (P < 0.02) (Figure 7H,I). We tested for differences in mRNA levels of selected candidate ChREBP and PPAR’Y target genes in liver (Figure 7J). This analysis identified increases in AZD1656 treated *Gckr*^del/wt^ mice for ChREBP-β and *Fgf21* a hepatokine that regulates peripheral lipolysis [39], and *Cidea* and *Cidec (Fsp27)* which encode lipid droplet and perilipin associated proteins involved in the regulation of lipid droplet dynamics, fusion and growth [40]. These changes implicate mechanisms linked to the raised circulating fatty acid concentration and hepatocyte steatosis in the AZD1656 treated *Gckr*^del/wt^ mice. Cumulatively the AZD1656 treated *Gckr*^del/wt^ mice had modestly raised glucose intolerance, raised blood cholesterol and free fatty acids and also increased liver triglyceride and hepatocyte microvesicular steatosis with a modest increase in sub-stage fibrosis. It is noteworthy that the mice of the *Gckr*^del/wt^ study were older and more insulin intolerant (P < 0.001) than the mice of the P446L study (Figure 8).

**Figure-8:**
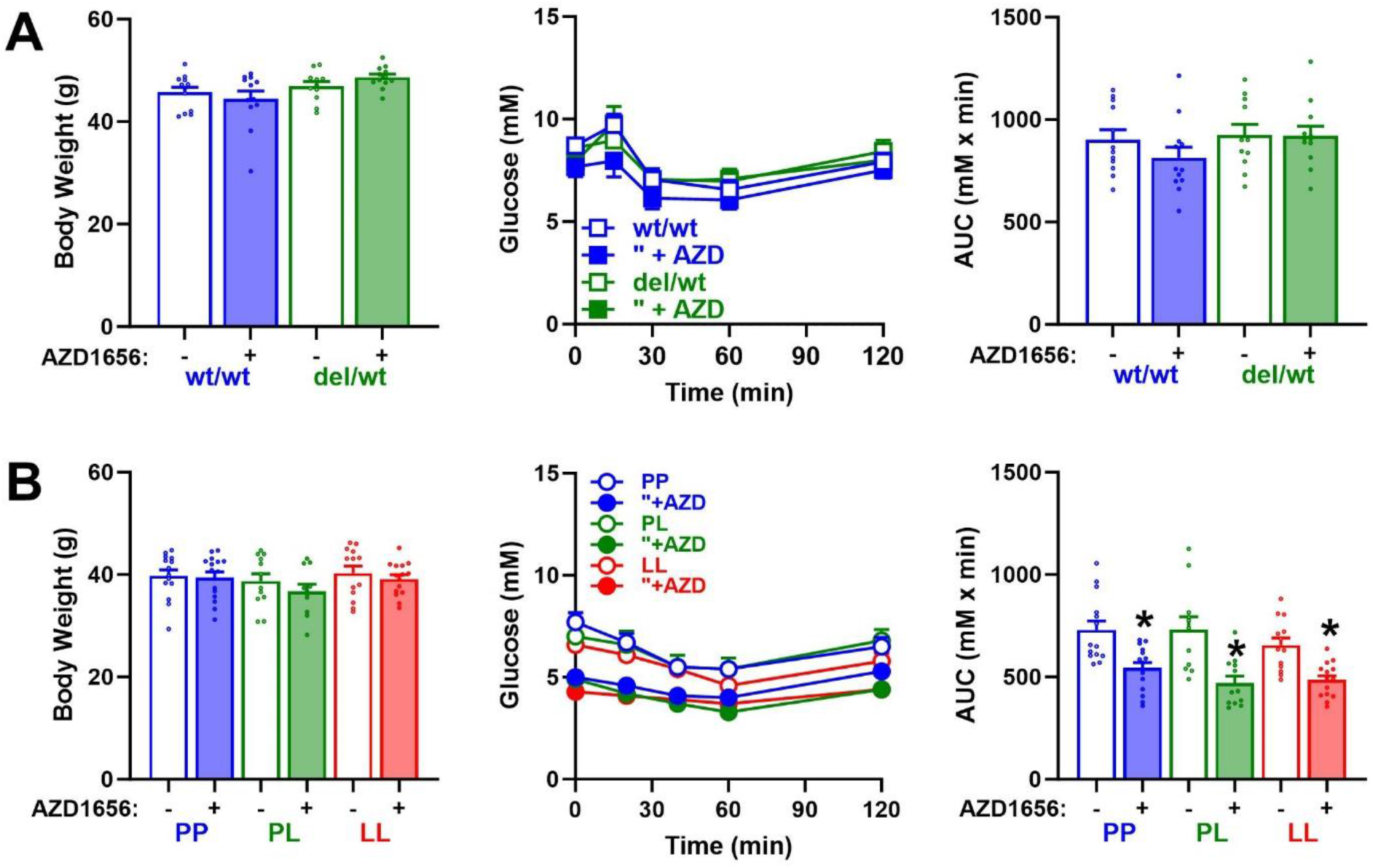
Insulin tolerance test in *Gckr*^del/wt^ and *Gckr:*P446L studies. A) *Gckr*^del/wt^ study: body weight and insulin tolerance test, blood glucose after insulin challenge and area under the curve (AUC) performed after 12-wk of HFD -/+ AZD1656 (3 mg/kg body wt) in mice aged 30 wk. B) *Gckr:*P446L study: body weight and insulin tolerance test performed after 12 wk of HFHSD -/+ AZD1656 (3 mg/kg body wt) in mice were aged 20 wk. * *P < 0*.*05* effect of AZD1656.

## 6. DISCUSSION

GKAs were developed for T2D and trialled as monotherapy [4,12] or combined with metformin, [5,11]. Whether the chronic decline in GKA efficacy is dependent on patient cohort, duration of diabetes or intrinsic GK activity in either islet β-cells or liver remains unknown [9,13,14]. The rationale for targeting GK activation for lowering blood glucose was in part based on evidence of compromised liver and islet GK in advanced T2D [2]. However raised GK activity in diabetes has also been reported [41,42]. T2D is a complex disease with variable disease progression and heterogeneous response to specific anti-hyperglycaemic therapies [43]. In this study we used two genetic mouse models to test the hypothesis that compromised GKRP function (P446L) or expression (Gckr^del/wt^) as occurs with common *GCKR* gene exonic and intronic variants in human populations [34,35,44] affects the response to GKA therapy. The results show that in both mouse models the chronic response to AZD1656 is genotype-dependent with declining glucose-lowering efficacy and increased hepatic steatosis being more prominent in the GKRP-deficient genotype.

In the Gckr-P446L mouse which models the human *GCKR* exonic rs1260326 variant [35], the blood glucose lowering efficacy of AZD1656 was sustained in the wild-type (PP) but declined in LL mice despite similar initial glycaemic efficacy that was independent of genotype. In the *Gckr*^del/wt^ model, which represents more modest GKRP deficiency than the LL mouse, GKA treatment was associated with modestly increased glucose intolerance and raised liver triglycerides and hepatocyte microvesicular steatosis in a GKRP-deficient manner. Accordingly, GKRP deficiency predisposed to declining GKA glycaemic efficacy in the P446L model and to increased hepatic lipids and glucose intolerance in the*Gckr*^del/wt^ model. In previous pre-clinical studies sustained GKA efficacy without accumulation of liver lipids had been reported in various rodent models [13], but not in the *db/db* mouse which showed loss in GKA glycaemic efficacy [22-24] and increased hepatic steatosis [23,24,45]. The two GKRP-deficient models used in this study represent functional GK excess, evident from greater relative GK distribution to the cytoplasm despite compensatory lower total GK protein and activity. They represent a more modest phenotype than the hyperglycaemic *db/db* mouse, which is characterized by severe insulin resistance and elevated liver and pancreatic GK activity [26,46]. The declining GKA efficacy was not associated with a decrease in liver GK protein indicating that the repression of the *Gck* gene that occurs in hepatocytes cultured with GKAs [47,48] does not manifest in vivo, possibly because of compensatory changes in insulin and glucagon secretion which are strong transcriptional regulators of the *Gck* gene [29]. Elevated liver GK activity after chronic treatment with dual-targeting GKAs concurs with previous findings [21,49].

Two key differences between the *Gckr*-P446L (HFHSD) study and the *Gckr*^del/wt^ (HFD) study are the lower plasma insulin in P446L mice (LL, LP lower than wild-type PP) compared with the higher insulin in *Gckr*^del/wt^ mice (del/wt higher than wt/wt) and the greater insulin intolerance in the *Gckr*^del/wt^ study compared with the P446L study (Figure-8). Whether these differences are intrinsic to these genetic deletion and knock-in models or are linked to the older age of the *Gckr*^del/wt^ study or the different diets (HFD and HFSHD) cannot be ascertained. Previous studies on other Gckr-deletion models found no significant difference in insulin levels on standard rodent diet but higher insulin on a high energy diet [50,51]. The lower insulin levels in LL mice relative to corresponding wild-type is one of several metabolic traits that is shared between this mouse model [35] and the genome-wide associated traits with the human *GCKR* rs1260326^P446L^ variant [34]. It is likely that the P446L model represents greater functional GK-excess than the Gckr-del model as supported by the smaller fractional decrease in GK-activity for PL compared with PP (< 15%) than for *Gckr*^del/wt^ compared with *Gckr*^wt/wt^ (∼30%). In *Gck*-transgenic models increased hepatic steatosis was found in conjunction with raised insulin in aged mice [52] but not in young liver-selective *Gck*-transgenic mice that were resistant to diet-induced hyperinsulinemia [53,54]. The hepatic steatosis in the GKA-treated *Gckr*^del/wt^ but not in GKA-treated P446L mice may be linked to the higher plasma insulin and/or greater insulin intolerance. A recent study on the insulin resistant Zucker rat showed that continuous exposure to AZD1656 during the fasting but not during the postabsorptive phase increased hepatic steatosis suggesting that predisposition to steatosis may be linked to the endocrine state [17]. The present findings of increased steatosis in the *Gckr*^del/wt^ mouse concur with a mechanism for increased steatosis during compromised regulation of liver GK by GKRP and additionally highlight a potential permissive role for insulin resistance. It is noteworthy that in human populations the association of *GCKR* variants with raised blood triglycerides or hepatic steatosis was more prominent in T2D with poor glycaemic control and in obesity-associated insulin resistance [55-57].

The decline in blood glucose lowering efficacy of AZD1656 in the P446L model in the absence of hepatic steatosis implicates other hepatic or pancreatic adaptations. The liver transcriptome analysis showed that the P446L genotype had a greater effect on liver gene transcripts than the GKA treatment in the wild-type mice, which were qualitatively similar but more modest, indicating that treatment with AZD1656 at a dose eliciting ∼ 3mM decrease in blood glucose represents a more modest functional liver GK excess than occurs with the P446L substitution. Remarkably, the GKA treatment in the LL genotype did not mimic the transcriptome changes that occurred in the wild-type mice indicating that the hepatic transcriptome adaptations to relative GK excess in the P446L genotype was maximally attained. This could in part explain the decline in glycaemic efficacy of the GKA in the LL genotype in comparison with wild-type. Although, there was no evidence for a decline in pancreatic islet GK immunostaining, changes in pancreatic islet function in the α-cells or β-cells cannot be excluded. An increase in second phase insulin secretion in man by dual targeting GKAs [58] that also increase liver GK protein in rodent models [49] has been reported.

The findings of this study demonstrating dependence of chronic GKA efficacy on GKRP deficiency are based on mouse models for the human exonic P446L variant [34,35] and transcriptional *Gckr* deficiency [44]. To determine whether these findings translate to man, will require genotyping of participants of GKA clinical trials and analysis of *GCKR* variant alleles relative to the trial endpoint data for blood triglyceride, cholesterol and liver enzymes [4,5,11,12]. The current findings have two potential implications relevant for GK targeting therapies for metabolic disease. First, for GKA activation therapy for diabetes for other GKAs in development [9] people who are homozygous for *GCKR* variants are predicted to be more likely to show adverse effects on lipids and loss in glycaemic efficacy. Second, recent studies on the decline in islet β-cell function in the *db/db* mouse [27] and mouse models of hyperglycaemia due to altered K-ATP channel function [59,60] have argued in support of GK inhibition rather than activation therapy for protecting against β-cell failure [25,60-62]. The present findings of predisposition to hepatic steatosis and declining GKA efficacy in GKRP-deficient models suggest that GK inhibition therapy could protect from hepatic steatosis in people with *GCKR* variants particularly in conjunction with hyperinsulinaemia analogously to the dyslipidaemia in association with *GCKR* variants which manifest more prominently in insulin resistance and poorly controlled diabetes [55,56].

## ABBREVIATIONS

AUC: area under the curve
AZD1656: a glucokinase activator (cas no. 919783-22-5) developed by AstraZeneca
C-H: cytoplasmic H-score
DEG: differentially expressed genes
del/del: homozygous *Gckr* knock-out mice
del/wt: heterozygous
*Gckr*: knock-out mice
*GCK*: glucokinase gene
*GCKR*: glucokinase regulatory protein gene
GK: glucokinase
GKA: glucokinase activator
GKRP: glucokinase regulatory protein
H & E: haematoxylin and eosin
H-score: [% weak] + [2 x % medium] + [3 x % high]
HFD: high-fat diet
HFHSD: high-fat high-sugar diet
LChol: liver cholesterol
LTG: liver triglycerides
LWt: liver weight
LL: homozygous *Gckr:*P446L knock-in mice
PL: heterozygous *Gckr:*P446L knock-in mice
PP: wild-type *Gckr:*P446L knock-in mice
P446L: *Gckr:*P446L knock-in mice
N-H: nuclear H-score
RD: regular diet
rs1260326: single nucleotide polymorphism (P446L) of *GCKR* gene
T2D: type 2 diabetes
TChol: total blood cholesterol
TG: triglycerides
wk: week
wt/wt: wild-type *Gckr* knock-out mice

## AUTHOR CONTRIBUTIONS

B.E.F. and S.S.C. designed and performed experiments, analyzed the data and co-wrote the paper. D.T. conducted the histopathology analysis and interpretation; A.A. and F.O. contributed to data acquisition. R.J.F., D.M.S., contributed to experimental design, data collection and analysis. All authors reviewed, discussed and approved the manuscript. L.A. directed the project, conceived and designed the experiments and co-wrote the paper. All authors approved the final version prior to submission.

## ACKNOWLEDGEMENTS

We thank Rainie Cameron for her help with the animal work, Laura Wilson for scanning of the IHC slides, Ann Hedley and Simon J. Cockell from the Newcastle University Bioinformatics Support Unit for their help and guidance with the RNA-seq analysis.

## DATA RESOURCE AND AVAILABILITY

RNA-seq data was deposited in the NCBI Gene Expression Omnibus (GEO) database under accession series GSE228698 <https://www.ncbi.nlm.nih.gov/geo/query/acc.cgi?acc=GSE228698>. Raw figure data is deposited at data.ncl under the collection 7065890

<https://doi.org/10.25405/data.ncl.c.7065890>. Individual figure DOIs are:

Figure-1

<u>10.25405/data.ncl.25189130</u>

Figure-2

<u>10.25405/data.ncl.25189199</u>

Figure-3

<u>10.25405/data.ncl.25189169</u>

Figure-4

<u>10.25405/data.ncl.25189181</u>

Figure-5

<u>10.25405/data.ncl.25189187</u>

Figure-6

<u>10.25405/data.ncl.25189193</u>

Figure-7

<u>10.25405/data.ncl.25189196</u>

Figure-8

<u>10.25405/data.ncl.25189208</u>

